# The trade-off between grain weight and grain number in wheat is explained by the overlapping of the key phases determining these major yield components

**DOI:** 10.1101/2024.02.28.582521

**Authors:** Lucas Vicentin, Javier Canales, Daniel F. Calderini

**Affiliations:** Graduate School, Faculty of Agricultural Science, Universidad Austral de Chile, Campus Isla Teja, Valdivia, Chile; Institute of Plant Production and Protection, Universidad Austral de Chile, Campus Isla Teja, Valdivia, Chile; Institute of Biochemistry and Microbiology, Universidad Austral de Chile, Campus Isla Teja, Valdivia, Chile; ANID-Millennium Science Initiative Program-Millennium Institute for Integrative Biology (iBio), Santiago, Chile

**Keywords:** TaExpA6, TaGW2, source-sink, expansins, ovary weight, grain size regulation, grain yield, cereals

## Abstract

Enhancing grain yield is a primary goal in the cultivation of major staple crops, including wheat. Recent research has focused on identifying the physiological and molecular factors that influence grain weight, a critical determinant of crop yield. However, a bottleneck has arisen due to the trade-off between grain weight and grain number, whose underlying causes remain elusive. In a novel approach, a wheat expansin gene, TaExpA6, known for its expression in root tissues, was engineered to express in the grains of the spring wheat cultivar Fielder. This modification led to increases in both grain weight and yield without adversely affecting grain number. Conversely, a triple mutant line targeting the gene TaGW2, a known negative regulator of grain weight, resulted in increased grain weight but decreased grain number, potentially offsetting yield gains. This study aimed to evaluate four wheat genotypes: (i) a transgenic line expressing TaExpA6, (ii) its wild-type counterpart (Fielder), (iii) a TaGW2 triple mutant line, and (iv) its wild-type. Conducted in southern Chile, the study employed a Complete Randomized Block Design with four replications, under well-managed field conditions including fertilization, irrigation, and pest control. The primary metrics assessed were grain yield, grain number, and average grain weight per spike, along with detailed measurements of grain weight and dimensions across the spike, and ovary weight at pollination (Waddington’s scale 10). The expression levels of TaExpA6 and TaGW2 were also monitored post-anthesis. Results indicated that both the TaExpA6 line and the triple mutant line achieved significantly higher average grain weights compared to their respective wild types. Notably, the TaExpA6 line did not exhibit a reduction in grain number, thereby enhancing grain yield per spike. In contrast, the triple mutant line showed a reduced grain number per spike, with no significant change in overall yield. Analysis of ovary size, grain weight dynamics, and gene expression patterns suggests that the trade-off between grain weight and number could be attributed to the overlapping of the critical periods for the determination of these traits.

## 1 Introduction

Advancements in the understanding of physiological and molecular control of grain weight and size in wheat and other staple food crops have been significant over the past decade (Brinton and Uauy, 2019; Slafer et al., 2023). This progress has been driven by rapid molecular developments, leading to the identification of key genes, transcription factors, and Quantitative Trait Loci (QTL) linked to grain weight and size in various crops (barley: Yang et al., 2023; rapeseed: Canales et al., 2021; rice: Zuo and Li, 2014; sorghum: Tao et al., 2021; sunflower: Castillo et al., 2018; wheat: Mira et al., 2023; Khan et al., 2022; Tillett et al., 2022; Adamski et al., 2021; Kumar et al., 2016; Simmonds et al., 2016; Simmonds et al. 2014). These studies assume that grain weight, which is a key yield component, could lead to the yield increase required to meet the challenge of food security. However, the relationship between increased grain weight and overall yield improvement is complex due to the reported trade-off between grain weight and grain number in wheat and other crops. In wheat, grain weight improvement has been addressed through various strategies, from classical breeding focusing on increasing grain weight through recurrent selection (Wiersma et al., 2001), to molecular breeding techniques such as mutating the *Ta*GW2 gene, a known negative regulator of grain size and weight (Yang et al., 2012; Hong et al., 2014; Wang et al., 2018; Zhang et al., 2018). Additional methods include gene introgression, like *Ta*GSNE (Khan et al., 2022), and overexpression of genes such as *Ta*BG1 and *Ta*CYP78A5 (Milner et al., 2021; Guo et al., 2022). Although most of these attempts successfully increased grain weight of wheat, they failed to improve grain yield due to the trade-off between grain weight and number (Wiersma et al., 2001; Okamoto and Takumi, 2013; Brinton et al., 2017; Milner et al., 2021; Mora-Ramirez et al., 2021).

The causes of the trade-off between grain weight and number are not fully understood, despite extensive research (Wiersma et al., 2001; Sadras, 2007; Dwivedi et al., 2021; Fischer, 2022; Slafer et al., 2023). Initial hypotheses and few subsequent studies suggested that growing grains of wheat are limited by the source of assimilates during grain filling (Sinclair and Jamieson, 2006; Golan et al., 2019; Rosati and Benincasa, 2023). However, most of the research has demonstrated that wheat grains are not, or scarcely, limited by assimilate supply during post-anthesis (Slafer and Savin, 1994; Borrás et al., 2004; Fischer, 2008; Serrago et al., 2013; Slafer et al., 2021; Murchie et al., 2023), except under extreme source restriction in high-yielding environments (Beed et al., 2007; Sandaña et al., 2009; Alonso et al., 2018). An alternative explanation involves the increased proportion of smaller distal grains when grain number is increased through breeding or crop management, since distal grains are intrinsically smaller than proximal ones (Acreche and Slafer, 2006; Ferrante et al., 2015; Ferrante et al., 2017). However, this does not account for situations where grain weight improvements lead to an actual trade-off with grain number (e.g., Wiersma et al., 2001; Brinton et al., 2017; Wang et al., 2018). Notably, interventions that successfully increased grain weight in both proximal and distal grain positions within the spike often resulted in a reduced grain number per spike and area (Wiersma et al., 2001; Okamoto and Takumi, 2013; Brinton et al., 2017; Quintero et al., 2018; Wang et al., 2018; Zhai et al., 2018; Adamski et al., 2021; Milner et al., 2021), confirming a genuine trade-off between yield components. Furthermore, recent genomic studies indicate that many regions associated with grain number (GN) and grain weight (GW) coincide and have inverse phenotypic effects, suggesting a strong genetic basis for this trade-off (Xie and Sparkes, 2021).

Trade-offs between grain number subcomponents have been documented in wheat. For instance, a higher plant number correlates with a lower number of spikes per plant (Slafer et al., 2021). It is generally accepted these trade-offs are due to the feedback between grain number components, whose settings overlap during the crop cycle (Slafer et al., 2021). For a long time, this explanation left aside the trade-off between grain weight and grain number, as these yield components were thought to have minimal overlap. From this perspective, the critical period for grain number determination accounts for 20 days before and 10 days after anthesis (DAA) (Fischer, 1985; Savin and Slafer, 1991; Abbate et al., 1997), whereas grain weight determination occurs during the grain filling period. However, several evidence now suggest that potential grain weight is established between booting and early grain filling in wheat (Calderini et al., 1999; Calderini and Reynolds, 2000; Ugarte et al., 2007; Hasan et al., 2011; Simmonds et al., 2016; Parent et al., 2017). This process is strongly influenced by maternal tissues, which impose a physical upper limit on grain weight in wheat (Millet and Pinthus, 1980; Calderini and Reynolds, 2000; Xie et al., 2015; Yu et al., 2015; Brinton et al., 2017; Reale et al., 2017; Brinton and Uauy, 2019; Calderini et al., 2021), a phenomenon also observed under increased temperature conditions (Calderini et al., 1999; Ugarte et al., 2007; Kino et al., 2020). The relevance of maternal tissues in determining grain size is also evident in other grain crops, for instance barley (Scott, et al., 1983; Ugarte et al., 2007; Radchuk et al., 2011), sorghum (Yang et al., 2009) and sunflower (Lindström et al., 2006; Lindström et al., 2007; Rondanini et al., 2009; Castillo et al., 2017). Additionally, the relationship between fruit size and flower ovary size has been reported in berries and fruit trees (Kiwifruit: Lai et al., 1990; Cruz-Castillo, 1991; olive: Rosati et al., 2009; peach: Scorzal et al., 1991; strawberry: Handley and Dill, 2003). In wheat, various molecular strategies have been explored to increase grain weight.

However, only a few have been successful in improving this trait (e.g. Hong et al., 2014; Simmonds et al., 2016; Brinton et al., 2017; Wang et al., 2018; Adamski et al., 2021; Jablonski et al., 2021; Milner et al., 2021; Mora-Ramirez et al., 2021), and even fewer have managed to increase grain weight without a trade-off with grain number (Calderini et al., 2021; Guo et al., 2022). Remarkably, one such improvement was achieved under field conditions at farmer’s plant density rate, where grain weight increased by 12.3% and grain yield by 11.3%, without affecting grain number (Calderini et al., 2021). Against this background, the present study aims to deepen the understanding of the trade-off between grain weight and grain number in wheat by evaluating lines with and without this trade-off. For this assessment, wheat lines previously tested under agronomic conditions were selected. We included the triple mutant line for the *Ta*GW2 gene (Wang et al., 2018), which releases grain growth by breaking the negative control of *Ta*GW2 over grain weight but demonstrates a trade-off with grain number. In contrast, the selected genotype without a trade-off is a line with ectopic expression of the expansin gene *Ta*ExpA6, which is naturally expressed in wheat roots but was cloned with a promoter for expression in growing grains (Calderini et al., 2021). Expansins are small proteins crucial for plant cell growth, facilitating cell wall stress relaxation induced by turgor pressure (McQueen-Mason et al., 1992; Cosgrove, 2021;2023). The expression of different expansins during wheat grain growth has been documented in wheat (Lin et al., 2005; Lizana et al., 2010; Kino et al., 2020; Xie and Sparkes, 2021; Mira et al., 2023). Both the triple mutant and transgenic lines were evaluated alongside their respective wild types (WT).

The ectopic expression of *Ta*ExpA6 and its protein in growing grains of wheat was evident from 10 DAA on (Calderini et al., 2021). This led the authors to propose that the trade-off between grain weight and grain number is potentially associated with the overlapping of the critical window for these two traits determinations. Our study aims to elucidate this trade-off by evaluating two genetically distinct wheat genotypes, known for increased grain weight but differing in the trade-off between both major yield components. These genotypes, along with their respective wild types, will be examined under field conditions. By dissecting grain yield components, along with physiological and molecular characteristics at the spike level, the study seeks to minimize confounding variables and clarify the underlying mechanisms of the observed trade-off.

## 2 Materials and Methods

### 2.1 Field conditions and experimental setup

Two field experiments were conducted on a Typic Hapludand soil at the Universidad Austral de Chile’s Experimental Station (EEAA) in Valdivia (39°47’S, 73°14’W). The first experiment spanned the 2021-2022 growing season (referred to as Exp. 1), while the second was conducted in the 2022-2023 season (Exp. 2). To fulfil the proposed objective, four spring wheat cultivars were selected for both experiments: (i) *Ta*ExpA6, a transgenic line expressing the expansin gene *Ta*ExpA6 ectopically in grains, as described by Calderini et al. (2021); (ii) its segregant wild type cv. Fielder; (iii) *Ta*GW2, a triple knock-out mutant of *Ta*GW2 (referred to as "aabbdd" in Wang et al., 2018); and (iv) its segregant wild type. These genotypes were chosen due to their contrasting effects on the trade-off between grain weight (GW) and grain number (GN). Specifically, while line *Ta*GW2 exhibits reduced GN, line *Ta*ExpA6 improves GW without impacting GN.

The *Ta*ExpA6 line features overexpression of *Ta*ExpA6 (REFSEQ v.1.1:

*TraesCS4A02G034200*) in the endosperm, aleurone, and pericarp tissues of developing grains. This overexpression is controlled by the wheat *puroindoline-b* (*PinB*) gene promoter (REFSEQ v.1.1: *TraesCS7B02G431200*), as detailed by Gautier et al. (1994) and Digeon et al. (1999). The *Ta*ExpA6 line and its segregant WT were developed and kindly shared by Dr. Emma Wallington from the National Institute of Agricultural Botany (NIAB), UK (Calderini et al., 2021). In contrast, the *Ta*GW2 gene in line *Ta*GW2 harbors mutations leading to a truncated, non-functional protein, as reported by Simmonds et al. (2016) and Wang et al. (2018). These GW2 lines were generously provided by Prof. Cristóbal Uauy from John Innes Center, UK.

Exp. 1 was sown on September 21, 2021. Due to phenological differences observed between groups of lines in Exp. 1, sowing dates in Exp. 2 were staggered: GW2 lines, with longer crop cycles, were sown earlier on August 20, 2022, while the shorter-cycle ExpA6 lines were sown later on September 2, 2022. Both experiments followed a randomized complete block design with four replications. Each plot measured 2 m in length and 1.2 m in width, consisting of 9 rows with 0.15 m spacing, and was sown at a density of 300 plants per square meter.

Optimal agronomic management was employed for all plots to prevent biotic and abiotic stress. Fertilization at sowing included 150 kg N ha^-1^, 150 kg P2O5 ha^-1^, and 100 kg K2O ha^-1^. An additional 150 kg N ha^-1^ was applied at tillering. To address potential aluminum toxicity brought about by low soil pH, the experimental site was treated with 4 Tn ha^-1^ of CaCO3 one month prior to sowing. Pests and diseases were managed using chemical treatments as per manufacturer recommendations. Drip irrigation supplemented rainfall to avoid water stress throughout the crop cycle.

Meteorological data, including air temperature and incident photosynthetically active radiation (PAR), were recorded daily from sowing until harvest at the Austral Meteorological Station of EEAA (http://agromet.inia.cl/), located approximately 150 m from the experimental plots.

### 2.2 Crop sampling and measurements

Crop phenology was recorded twice weekly according to the decimal code scale (Zadoks et al., 1974). At harvest, 45 spikes of main stems were sampled along 1m from the central row of each plot in both experiments. Grain yield per spike, grain number per spike and average grain weight were measured or calculated as previously (Bustos et al., 2013; Quintero et al., 2018). In addition, 10 more main shoot spikes of similar development and size were sampled from each plot to quantify grain weight and dimensions (length and width) at each grain position from every spikelet of the spike. From each spike, half the spikelets (i.e. all the spikelets along one side of the spike) were measured, considering the spike symmetry. Grains from positions G1 to G4 (G1 being the closest grain to the rachis and G4 the most distal, if present) within each spikelet were taken out, oven dried (48 h at 65 °C) and weighted separately using an electronic balance (Mettler Toledo, XP205DR, Greifensee, Switzerland). The length and width of each grain was recorded using a Marvin Seed Analyzer (Marvitech GmbH, Wittenburg, Germany). Grain number per spike and per spikelet were also recorded.

In Exp. 2, the weight of ovaries from florets at positions G1 to G4 was measured at pollination (stage 10 in Waddington et al., 1983) by sampling 20 ovaries of each floret position from the four central spikelets of five spikes per plot. In this experiment the time-course of grain weight and dimensions were also measured. From anthesis onwards, four main shoot spikes were sampled from each experimental unit twice weekly until physiological maturity to record grain fresh and dry weight and dimensions (i.e. length and width) of six individual grains corresponding to a proximal (G2) and a distal (G3) grain position of two central spikelets. The fresh weight and the length and width of grains were recorded immediately after sampling as described above. Dry weight of grains was measured with the same electronic balance, after drying the samples at 65 °C in an oven for 48 h.

In both experiments, grain quality was assessed by measuring grain protein concentration to determine the impact of GW changes on this quality trait. Accordingly, grains from positions G1 to G3 within the four central spikelets of the 10 main shoot spikes samples were bulked at each experimental unit and then milled using a Perten 120 laboratory mill (Huddinge, Sweden). The quantification of total nitrogen was executed using the Kjeldahl method. Protein content was then calculated by multiplying the total nitrogen value by a factor of 5.7, in accordance with the approach of Merrill and Watt (1973) as applied in Lizana and Calderini (2013).

### 2.3 Time-course expression analyses by reverse transcription quantitative PCR

To elucidate the relationship between grain growth dynamics and gene expression, we conducted time-course expression analyses of *PinB*::*Ta*ExpA6 in the ExpA6 lines and *Ta*GW2 in the GW2 lines. These analyses were performed using reverse transcription quantitative PCR (RT-qPCR) on grains at positions G1 and G2. Specifically, G1 and G2 grains from central spikelets on the main stem spikes were collected from a minimum of eight spikes at 4, 7, 10, 14, and 21 days after anthesis (DAA) for each experimental plot.

Immediately following collection, samples were secured in cryotubes, snap-frozen in liquid nitrogen, and subsequently preserved at -80°C until further processing. Total RNA was isolated using NucleoSpin™ columns (Macherey-Nagel), employing a standardized protocol adapted from Sangha et al. (2010). RNA samples were then subjected to DNaseI (Invitrogen) treatment, and cDNA synthesis was performed using the ImProm-II™ Reverse Transcription System with an input of 500 ng RNA per reaction.

Quantitative PCR (qPCR) was carried out in a 25 μL reaction volume using the Brilliant II SYBR Green PCR Master Mix (Stratagene, Agilent technologies). Primer concentrations were set at 0.2 μM. For *Ta*ExpA6, the primers TransgeneTaExpa6_F1 (5’-ATCTCCACCACCACCAAAACA-3’) and TransgeneTaExpa6_R1 (5’-GAAGCAGAACGCGAGAACGG-3’) were used. In the case of *Ta*GW2, genome-specific primers for its homoeologues, as described by Wang et al. (2018), were utilized. Controls without template and transcriptase were included to check for genomic DNA contamination.

The relative mRNA abundance of the target genes in grain tissues was determined using the comparative CT (ΔΔCT) method, as proposed by Pfaffl (2001). The ubiquitin conjugating enzyme (*Traes_4AL_8CEA69D2E.1*) served as the internal reference gene, amplified with primers Traes_4AL_8CE_esF (5’-CGGGCCCGAAGAGAGTCT-3’) and Traes_4AL_8CE_esR (5’-ATTAACGAAACCAATCGACGGA-3’). Data analysis for gene expression quantification was conducted using LinRegPCR software (Ruijter et al., 2009).

### 2.4 Statistical analysis

Analysis of variance (ANOVA) was applied to evaluate the effect of genotype on main shoot grain yield and associated traits, by using Statgraphics Centurion 18 software.

Fisher’s least significant difference test post hoc and/or Student’s t-test were employed to identify each significant difference within the evaluated group of lines. Additionally, a Two-Way ANOVA analysis was performed to assess significant differences in *Ta*ExpA6 and *Ta*GW2 gene expression between modified lines and their respective WTs using GraphPad Prism 8 software. Linear regression analysis was performed to assess the associations between measured grain traits. A tri-linear model was fitted to estimate the rate and duration of the lag and linear phases of grain filling, and final grain weight in the individual seed weight dynamics. Extra sum of squares F test was used for the comparison of the slopes and timings between each modified line and its respective WT. To model the dynamics of individual grain water content during the grain filling period, a second order polynomial model was employed. All model fittings were performed using GraphPad Prism 8 software.

## 3 Results

### 3.1 Environmental conditions and crop phenology across seasons

Environmental conditions from seedling emergence to physiological maturity were consistent between experiments (Table 1). Temperature variations between Exps. 1 and 2 were minimal, with differences of less than 1°C observed during the emergence-anthesis and grain filling periods. Incident PAR differed between the experiments, being 2.3 MJ m^-2^ d^-1^ higher in Exp. 1 than in Exp. 2 during the Emergence-Booting (Em-Bo) and grain filling phases (10.4 vs. 8.1 MJ m^-2^ d^-1^ and 12.8 vs. 10.5 MJ m^-2^ d^-1^, respectively). However, no significant difference in PAR was noted during the Booting-Anthesis (Bo-An) period.

**Table 1.** Average mean temperature, incident photosynthetically active radiation (PAR), photothermal quotient (Q) and photoperiod during the Emergence-Booting (Em-Bo), Booting-Anthesis (Bo-An) and Anthesis-Physiological Maturity (An-PM) periods in experiments 1 and 2.

| Experiment | Line* | Mean temperatures (°C) |  |  | Incident PAR (MJ m<sup>-2</sup> d<sup>-1</sup>) |  |  | Q (MJ m<sup>-2</sup> d<sup>-1</sup> °C<sup>-1</sup>)** |  |  | Photoperiod (h) |
| --- | --- | --- | --- | --- | --- | --- | --- | --- | --- | --- | --- |
|  |  | Em - Bo | Bo - An | An - PM | Em - Bo | Bo - An | An - PM | Em - Bo | Bo - An | An - PM | Em - An |
| Exp.1 (2021-2022) | TaExpA6 / WT<sub>TaExpA6</sub> | 12.3 | 14.2 | 17.2 | 10.3 | 12.3 | 12.8 | 2.9 | 2.6 | 2.1 | 14.6 |
|  | TaGW2 / WT<sub>TaGW2</sub> | 12.9 | 16.4 | 16.7 | 10.6 | 12.2 | 12.8 | 2.8 | 2.2 | 2.2 | 14.8 |
| Exp. 2 (2022-2023) | TaExpA6 / WT<sub>TaExpA6</sub> | 11.2 | 14.5 | 16.1 | 7.9 | 12.1 | 10.8 | 3.2 | 2.4 | 1.9 | 14.1 |
|  | TaGW2 / WT<sub>TaGW2</sub> | 11.3 | 15.4 | 16.3 | 8.2 | 13.2 | 10.3 | 3.1 | 2.5 | 1.8 | 14.1 |

\* Since no phenological difference was recorded between modified lines and their respective WT, values were calculated for the mean duration of each phenological period for the ExpA6 and GW2 line groups. \*\* Photothermal quotient was calculated as the ratio of mean daily incident radiation to mean temperature above 4.5 °C

Despite these variations, the photothermal quotient (Q) remained constant at 2.5 MJ m^-2^ d^-1^ °C^-1^ across the entire crop cycle in both seasons. A comparative analysis of the ExpA6 and GW2 genotype lines revealed similar weather exposure in both experiments, except for the Bo-An period in Exp. 1, where GW2 lines experienced a mean temperature 2.2°C higher than ExpA6 lines, leading to a 17% lower photothermal quotient for GW2 lines during this phase.

The length of the crop cycle averaged 123 days for ExpA6 lines and 142 days for GW2 lines across experiments (Fig. 1). In experiments 1 and 2, GW2 lines reached physiological maturity 13 and 26 days later, respectively, than ExpA6 lines. This delay is primarily attributed to the longer Emergence-Booting (Em-Bo) period in GW2 lines, as they exhibited a slower development rate. Post-booting, the differences in phenological stages, specifically Booting-Anthesis (Bo-An) and Anthesis-Physiological Maturity (An-PM), were negligible (i.e. less than 2 days) between the line groups. Despite minor climatic differences, the crop cycle in Exp. 2 extended by 16 and 27 days for ExpA6 and GW2 lines, respectively, compared to Exp. 1. This extension corresponds to an accumulated difference of 84 and 180 °Cd for each line group (Fig. S1). The extended crop cycle in GW2 lines, particularly in Exp. 2, is likely due to their heightened sensitivity to photoperiod compared to ExpA6 lines (see photoperiods in Table 1).

**Figure 1.**
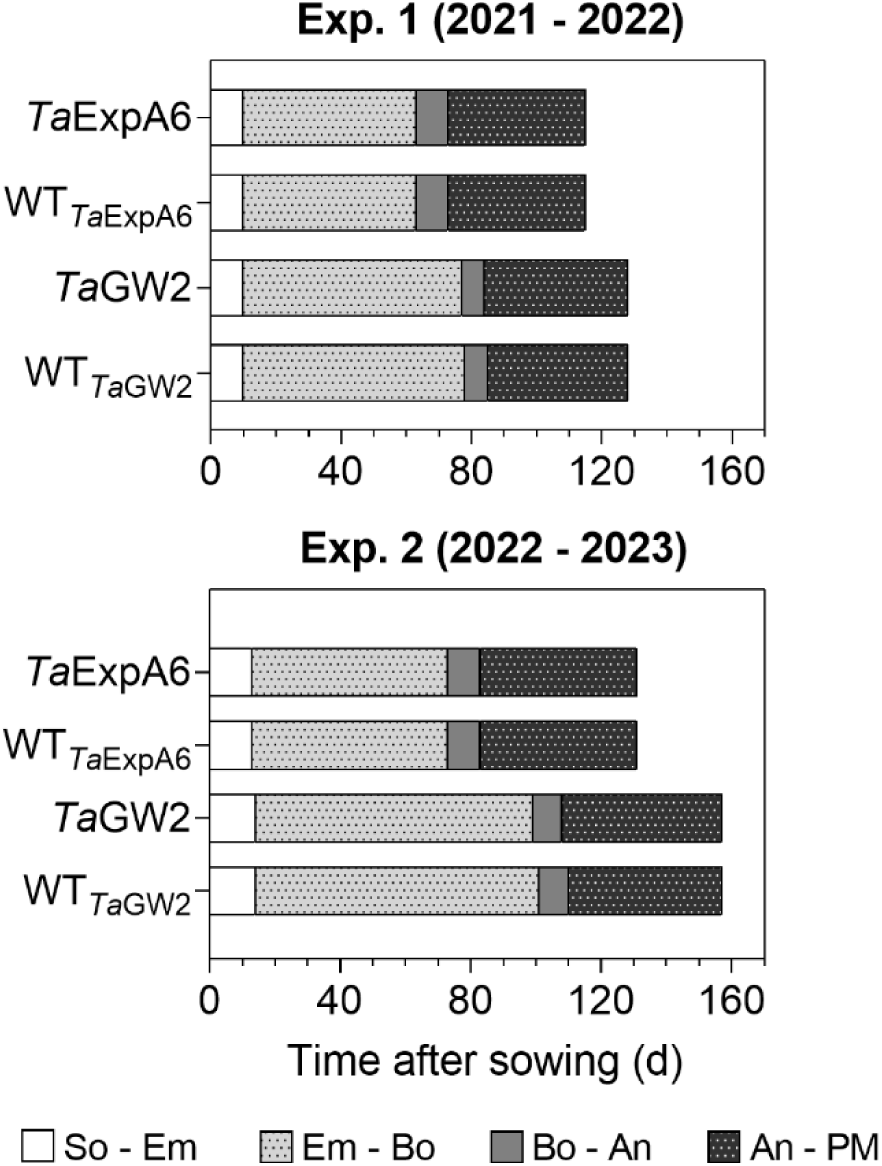
Phenological phases of *Ta*ExpA6 and *Ta*GW2 lines, and their respective wild types (WT), from sowing to physiological maturity in experiments 1 and 2. Bars show the duration of each phase in days: from sowing to seedling emergence (So-Em), from seedling emergence to booting (Em-Bo), from booting to anthesis (Bo-An) and from anthesis to physiological maturity (An-PM).

### 3.2 Grain yield per spike, yield components per spike and quality trait

Differences in grain yield per spike were observed between line groups (P < 0.05), with GW2 lines exhibiting higher yields compared to ExpA6 lines in both experimental seasons, averaging 2.39 g and 1.94 g, respectively (Table 2). Notably, the ectopic expression of *Ta*ExpA6 gene resulted in a significant increase in spike yield compared to the wild type (WT), showing increments of 14.0% and 8.2% in Exps. 1 and 2, respectively (P < 0.05). In contrast, the triple mutant of *Ta*GW2 and its WT displayed similar spike yields in both experiments (P < 0.05) (Table 2).

**Table 2.** Grain yield per spike (GY Spike^-1^), grain number per spike (GN Spike^-1^), average grain weight (TGW) and protein concentration (%) of grains recorded in *Ta*ExpA6 and *Ta*GW2 lines and their WTs in the field experiments 1 and 2.

| Experiment | Genotype (G) | Transgenesis or Tilling (T) | Main Stem Spike |  |  |  |  |  |  |  |
| --- | --- | --- | --- | --- | --- | --- | --- | --- | --- | --- |
|  |  |  | GY Spike<sup>-1</sup> (g) |  | GN Spike<sup>-1</sup> |  | TGW (g) |  | Protein (%) |  |
|  |  |  | Mean | s.e.m | Mean | s.e.m | Mean | s.e.m | Mean | s.e.m |
| Exp. 1 | ExpA6 lines | *TaExpA6* | 2.12 a | 0.06 | 42.4 a | 1.0 | 50.0 a | 1.1 | 11.1 a | 0.3 |
|  |  | WT<sub>*TaExpA6*</sub> | 1.86 b | 0.03 | 40.4 a | 0.3 | 46.0 b | 1.0 | 10.9 a | 0.5 |
|  | GW2 lines | *TaGW2* | 2.33 a | 0.10 | 43.6 b | 1.6 | 53.4 a | 1.0 | 11.5 a | 0.3 |
|  |  | WT<sub>*TaGW2*</sub> | 2.18 a | 0.10 | 50.4 a | 1.5 | 43.3 b | 0.8 | 10.4 b | 0.2 |
|  | ANOVA p-value (G) |  | * |  | ** |  | ns |  | ns |  |
|  | ANOVA p-value (T) |  | * |  | ns |  | ** |  | ns |  |
|  | ANOVA p-value (G*T) |  | ns |  | * |  | ** |  | ns |  |
|  | *t*-test p-value (*TaExpA6*) |  | ** |  | ns |  | * |  | ns |  |
|  | *t*-test p-value (*TaGW2*) |  | ns |  | * |  | ** |  | * |  |
| Exp. 2 | ExpA6 lines | *TaExpA6* | 1.97 a | 0.03 | 39.4 a | 0.4 | 50.0 a | 0.5 | 10.6 a | 0.1 |
|  |  | WT<sub>*TaExpA6*</sub> | 1.82 b | 0.04 | 39.6 a | 0.7 | 46.1 b | 0.4 | 11.2 a | 0.3 |
|  | GW2 lines | *TaGW2* | 2.68 a | 0.09 | 47.1 b | 1.0 | 56.7 a | 0.9 | 11.4 a | 0.2 |
|  |  | WT<sub>*TaGW2*</sub> | 2.38 a | 0.11 | 51.8 a | 1.2 | 45.7 b | 1.1 | 10.4 a | 0.6 |
|  | ANOVA p-value (G) |  | ** |  | ** |  | ** |  | ns |  |
|  | ANOVA p-value (T) |  | * |  | * |  | ** |  | ns |  |
|  | ANOVA p-value (G*T) |  | ns |  | * |  | ** |  | * |  |
|  | *t*-test p-value (*TaExpA6*) |  | * |  | ns |  | ** |  | ns |  |
|  | *t*-test p-value (*TaGW2*) |  | ns |  | * |  | ** |  | ns |  |

ANOVA P-value is shown in the table. All data are shown as mean and SEM. The phenotype data of each transformed/mutant line was compared with the respective WT using Student's *t*-test; different letters indicate significant effects: \*, *P* < 0.05; \*\*, *P* < 0.01; ns, not significant.

Regarding grain yield components, genotype significantly influenced these traits, contingent on the line group (P < 0.05). Average grain weight (TGW) was similar between both groups in Exp. 1 (P > 0.05) but was marginally higher in the GW2 lines (6.6% increase, P < 0.01) in Exp. 2, with weights of 51.2 g and 48.1 g for GW2 and ExpA6 line groups, respectively (Table 2). Both *Ta*ExpA6 and *Ta*GW2 lines surpassed the grain weight of their respective WTs in both experiments (P < 0.001). Averaging across Exps. 1 and 2, *Ta*ExpA6 and *Ta*GW2 lines exhibited increases in TGW of 8.6% and 23.7% above their WTs, respectively (Table 2). Conversely, contrasting results were found for the grain number per spike; the *Ta*ExpA6 line maintained a similar number to its WT (P > 0.05), whereas the *Ta*GW2 line exhibited a reduction in grain number compared to its WT in both experiments, with declines of 13.5% and 9.1% in Exps. 1 and 2, respectively (Table 2, Fig. S2). These findings align with previous reports indicating no trade-off in the *Ta*ExpA6 overexpressed line and a detrimental effect of the *Ta*GW2 loss-of-function triple mutant on grain number.

In addition, we assessed a critical wheat quality trait, such as grain protein concentration. For this analysis, grains from G1 to G3 positions of the four central spikelets were pooled from each experimental unit. The results indicated no significant impact of transgenesis or tilling on grain protein concentration (P > 0.05), although an interaction (P < 0.05) between these factors was observed (Table 2). Grains from plants with the *Ta*ExpA6 gene exhibited similar protein concentrations to the WT, highlighting the relevance of the *Ta*ExpA6 construct. Conversely, the *Ta*GW2 triple mutant either increased (by 1.1 percentage points) or did not affect grain protein concentration in Experiments 1 and 2, respectively.

Furthermore, no correlation (R^2^ = 0.05; P > 0.05) was observed between grain protein concentration and grain yield per spike across genotypes and experiments, suggesting that the *Ta*GW2 triple mutation improved this quality trait independently from a dilution-concentration effect.

### 3.3 Individual grain weight and dimensions along the spike

To have a deeper understanding of the impact of *Ta*ExpA6 overexpression and the *Ta*GW2 triple mutation, we dissected the spike evaluating grain weight, number and dimensions at each grain position along the spike (usually referred to as spike map) in both experiments. Our analysis revealed that across spikelets and grain positions, *Ta*ExpA6 overexpression resulted in an increase in grain weight compared to its wild type (WT) in both experiments. Specifically, grain weight enhancement in the *Ta*ExpA6 line was observed as follows: in Exp.1, grains at positions G1, G2, G3, and G4 showed increases of 9%, 8%, 11%, and 25%, respectively, and in Exp. 2, increases of 9%, 6%, 10%, and 10% for G1, G2, G3, and G4, respectively (see Fig. 2). In comparison, the *Ta*GW2 triple mutant exhibited a more pronounced increase in grain weight above the WT across all grain positions in both experiments: in Exp. 1, increases of 19%, 21%, 30%, and 37% for G1, G2, G3, and G4, respectively, and in Exp. 2, increases of 23%, 23%, 18%, and 19% for the same positions (Fig. 2). On the other hand, when averaging across experiments, the ExpA6 lines showed similar grain number per spikelet across the spike (i.e. -1%), while the triple mutation of *Ta*GW2 gene caused a reduction of 11% in this trait in regard to its WT (Fig. S2).

**Figure 2.**
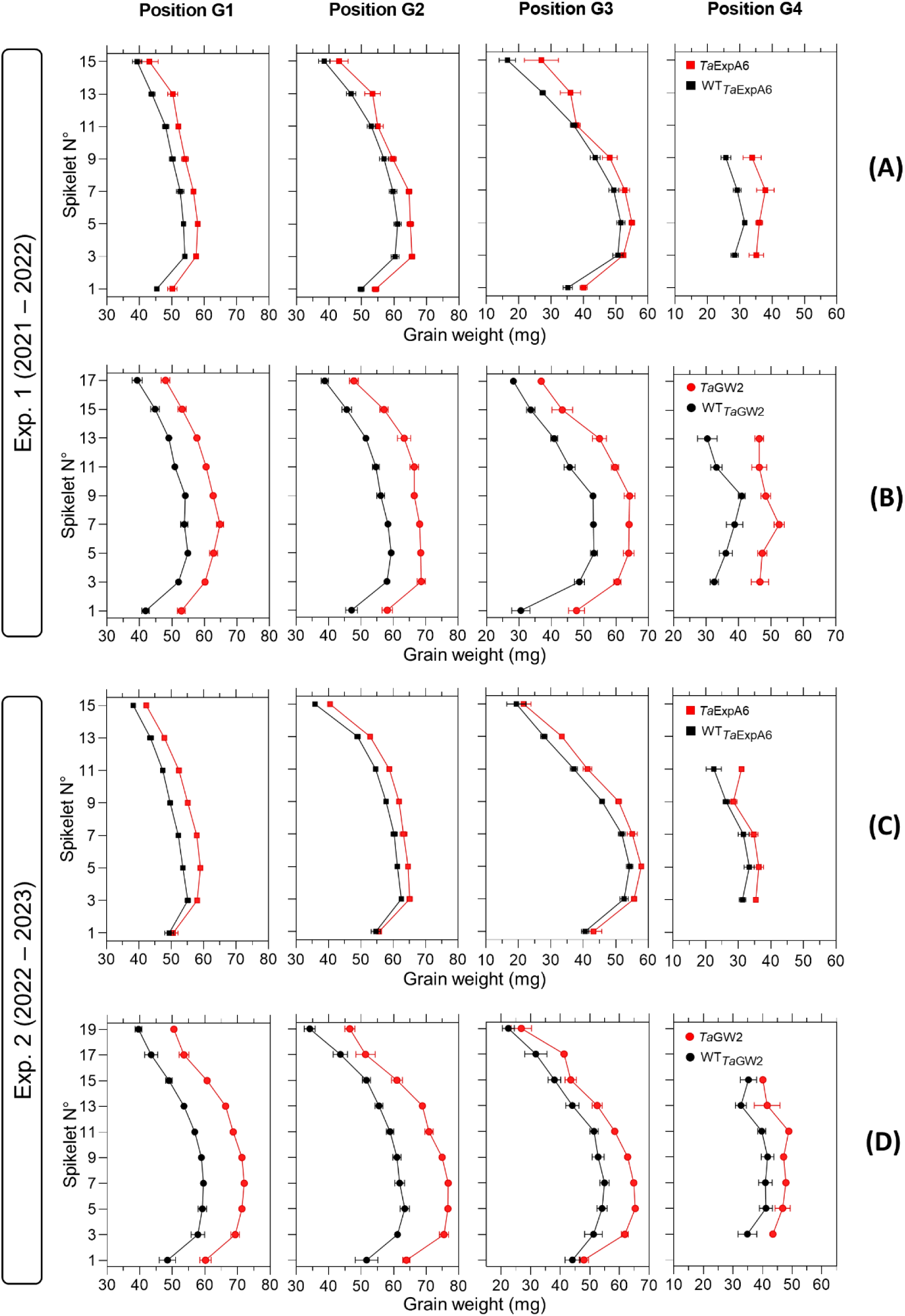
Grain weight in grain positions G1, G2, G3 and G4 from each spikelet along the spike of ExpA6 lines (A, C) and GW2 lines (B, D) in experiments 1 and 2. The transgenic *Ta*ExpA6 line and the triple mutant of *Ta*GW2 gene, and their WTs are depicted by red and black symbols, respectively.

Subsequent analysis focused on the relationship between grain dimensions and final grain weight. A strong association (P < 0.001) was observed between grain weight and both grain length and width, with determination coefficients ranging from 0.92 to 0.96 (Fig. 3). This relationship was consistent within each genotype group (Fig. S3).

**Figure 3.**
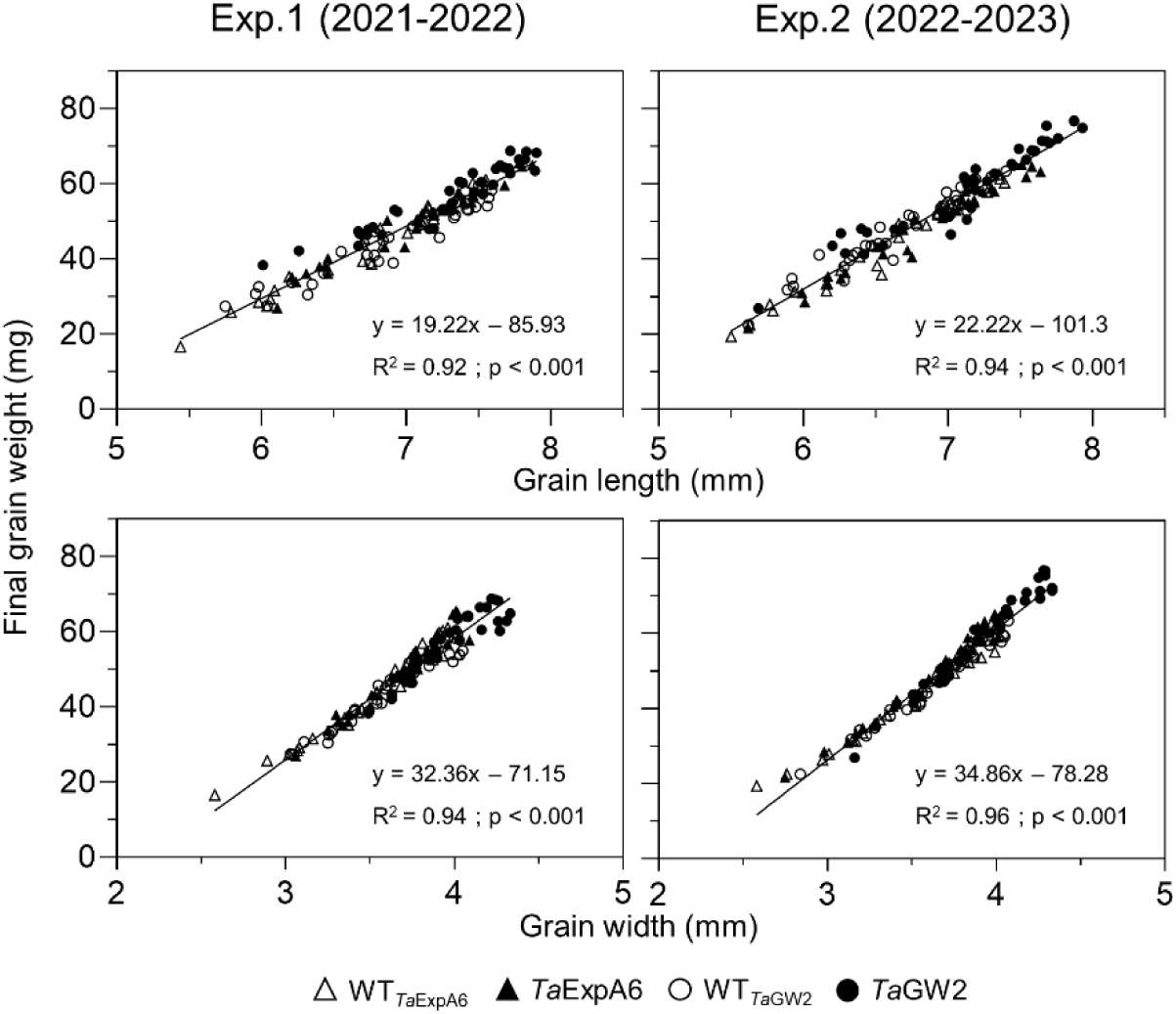
Relationship between grain weight and grain length (upper panel) or grain width (lower panel) of grain positions G1, G2, G3 and G4 from each spikelet along the spike, across genotype groups in experiments 1 (left panel) and 2 (right panel). *Ta*ExpA6 line and its WT are denoted by closed and open triangles respectively, while the *Ta*GW2 triple mutant line and its WT are denoted by closed and open circles, respectively.

### 3.4 Ovary weight, grain weight dynamics and gene expression

In Exp. 2, we assessed ovary weight at pollination (stage 10 according to Waddington et al., 1983) in florets at positions F1, F2, F3 and F4 from the central spikelets of spikes in each evaluated line. A linear association between final grain weight and ovary weight was found across lines and grain positions (Fig. 4). In agreement with this association, GW2 lines exhibited increased ovary weight compared to ExpA6 genotypes (Figs. 4 and 5). However, contrasting results were also found between both groups of lines, as the *Ta*GW2 triple mutant showed a significant increase (P < 0.05) in ovary weight compared to its WT in all but the G3 floret position, whereas the *Ta*ExpA6 construct showed no significant (P > 0.05) alteration in ovary weight at pollination, mirroring its WT (Fig. 5). This suggests that *Ta*GW2 gene mutation impacts ovary size prior to anthesis.

**Figure 4.**
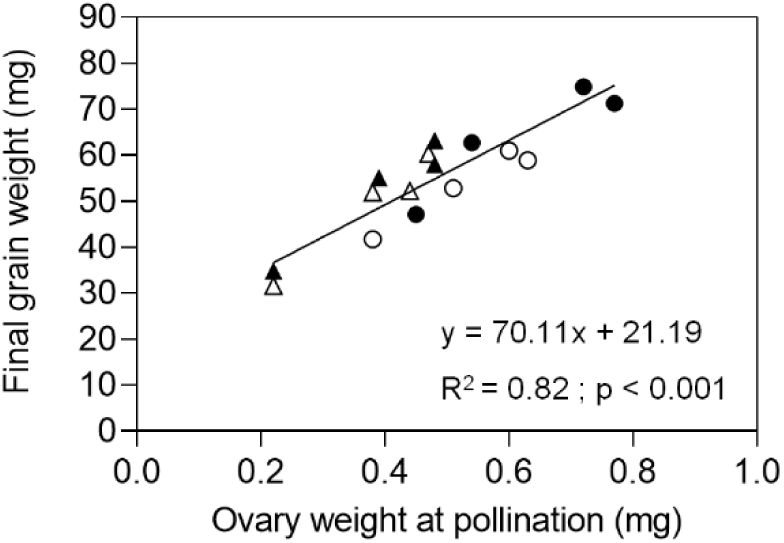
Relationship between final grain weight and ovary weight at pollination (W10, Waddington et al., 1983) of grain positions G1, G2, G3 and G4 from the central spikelets of the spike corresponding to the *Ta*ExpA6 line (closed triangles), *Ta*GW2 line (closed circles) and their WTs (open triangles and open circles, respectively).

**Figure 5.**
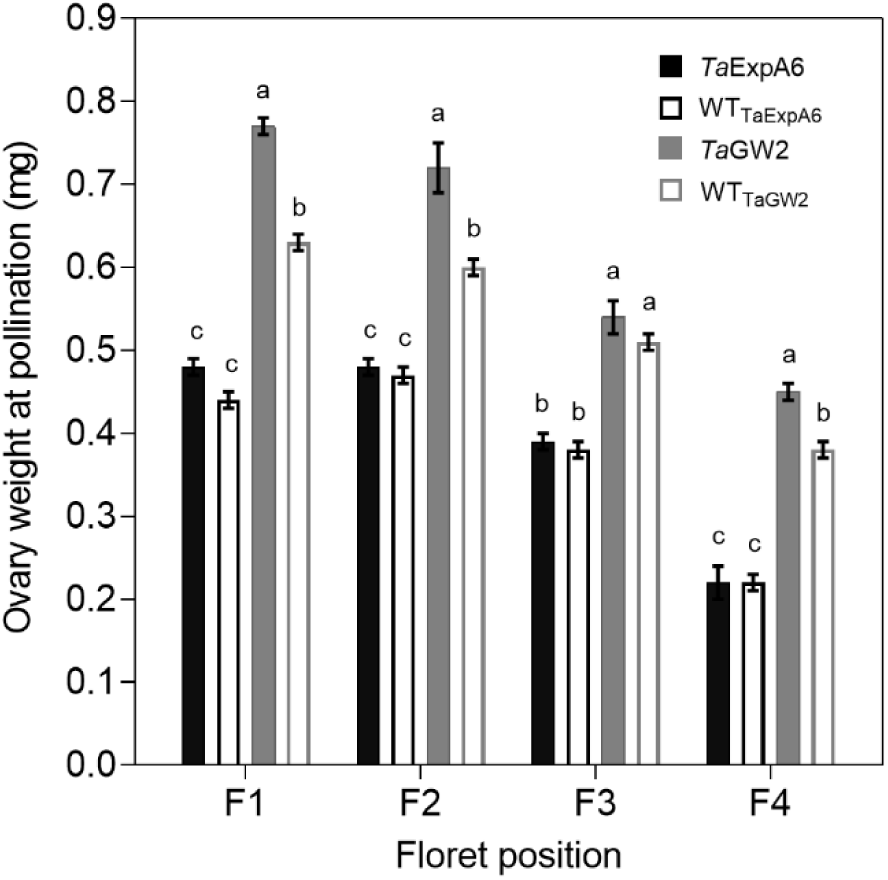
Ovary weight at pollination (W10, Waddington et al., 1983) of florets set at floret positions F1, F2, F3 and F4 from the central spikelets of the spike corresponding to the *Ta*ExpA6 line (solid black bars), *Ta*GW2 triple mutant line (solid grey bars) and their WTs (empty black and grey bars, respectively).

When the time-course of grain weight from positions G2 and G3 was monitored through the grain filling period, both modified lines (*Ta*ExpA6 and *Ta*GW2) surpassed their respective WTs in grain weight, though, the onset of these differences varied between groups. The GW2 triple mutant showed higher grain weight than the WT from the starting of measurements at 4 DAA (Fig. 6D, Fig. S4), while the *Ta*ExpA6 line exhibited higher grain weights at these grain positions from 20 DAA on (Fig. 6A, Fig S4). For both line groups, a tri-linear function accurately depicted individual grain weight dynamics (Fig. S5), with enhanced grain filling rates at the linear phase accounting for the increased final grain weights in both G2 and G3 grains (R^2^ = 0.99; P < 0.05), with no significant difference in the duration of grain filling, which was approximately 40 days (Table S1). Higher grain weights in modified lines were also coupled with an increased maximum grain water content (Figs. 6A and D; Fig. S4).

**Figure 6.**
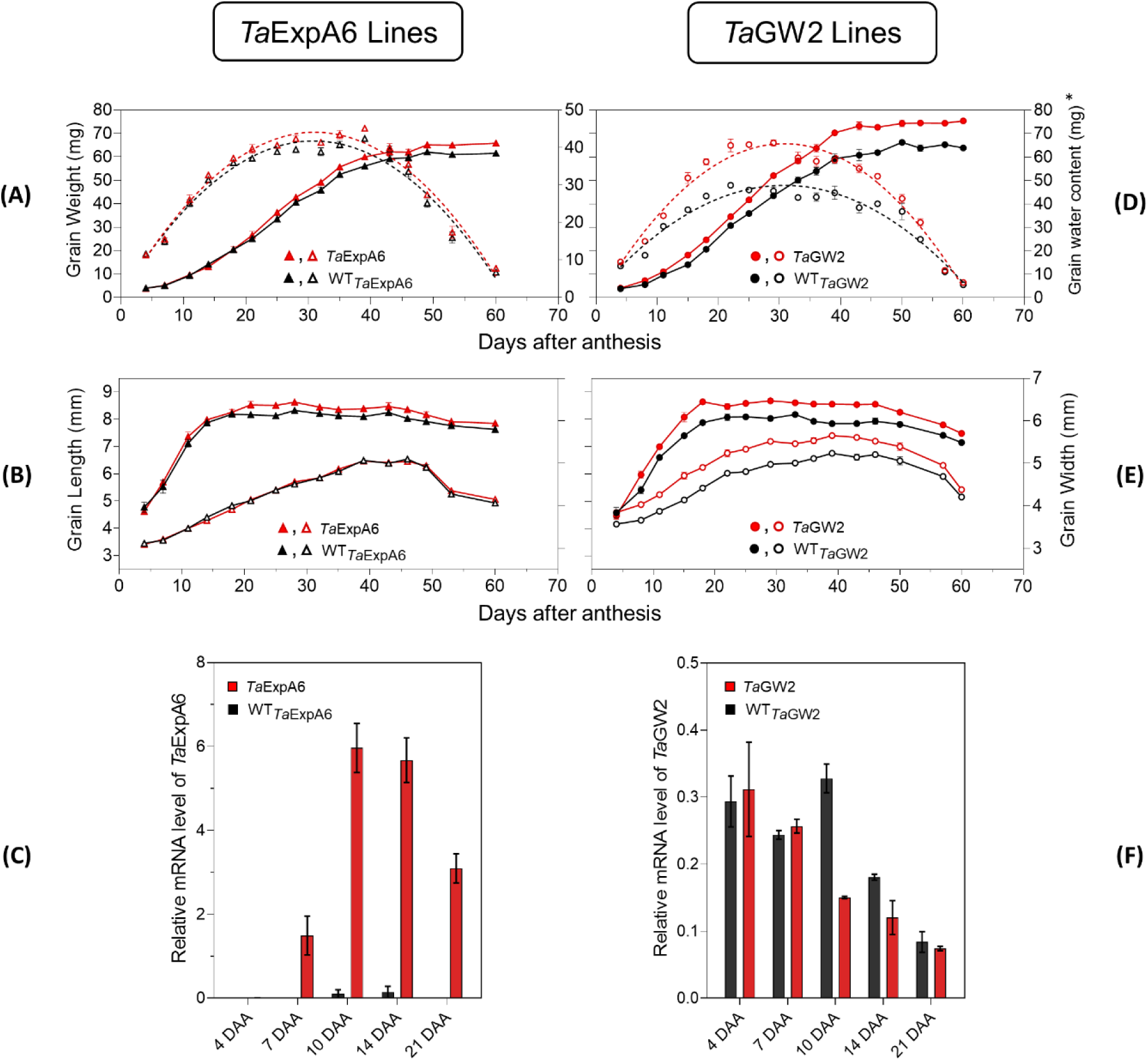
Grain weight dynamics and gene expression. (A) Grain weight and water content dynamics at grain position G2 from the central spikelets of the spike corresponding to the ExpA6 lines and (D) GW2 lines. (B) Grain length and grain width dynamics at the same grain position corresponding to the ExpA6 lines and (E) GW2 lines. (C) Relative *Ta*ExpA6 gene expression level in developing grains in the ExpA6 lines. (F) Relative *Ta*GW2 gene expression level in developing grains in the GW2 lines. In all cases, bars show the standard error of the means. *\*Note that different scales were used to plot grain water content of ExpA6 and GW2 lines*.

Grain dimension dynamics also varied between the line groups. The *Ta*ExpA6 line increased grain length by 3% and 4% in grain positions G2 and G3, respectively, over the WT (P < 0.05) without affecting grain width (Fig. 6B, Fig. S4). Conversely, the GW2 triple mutant improved both grain length and width along grain filling (P < 0.05) by 6.2 and 6.7%, respectively, when both grain positions were averaged (Fig. 6E, Fig. S4). Notably, differences in grain width in the ExpA6 lines became evident only at the ending of dimension dynamics analysis, coinciding with reduced grain water content (< 30%) (Fig. 6B, Fig. S4).

The observed divergence in grain length between the transgenic *Ta*ExpA6 line and its WT counterpart became apparent at 20 days after anthesis (DAA). This divergence appears to coincide with the expression profile of the *Ta*ExpA6 gene. Remarkably, *Ta*ExpA6 expression remains undetectable until 5 DAA, subsequently peaking between 10 and 15 DAA (see Figure 6C). In contrast, the grain dimensions in the GW2 triple mutant consistently exceeded those of the WT throughout the observation period (Fig. 6E).

Furthermore, *Ta*GW2 expression was monitored from 4 to 21 DAA, as depicted in Figure 6F. It is critical to note that the triple mutation in *Ta*GW2 induces alterations in both the DNA sequence and the resultant functional protein, without impacting gene expression levels (Wang et al., 2018). This is reflected in the two-way ANOVA results, showing modest genotypic effects for *Ta*GW2 (p = 0.0243) compared to the more significant effects in *Ta*ExpA6 (p < 0.0001). Furthermore, Bonferroni’s multiple comparisons test revealed significant changes in *Ta*GW2 only at 10 DAA (adjusted p-value < 0.05, Figure 6F). In contrast, *Ta*ExpA6 exhibits consistent and significant changes in mRNA levels at multiple time points, specifically at 10, 14, and 21 DAA (adjusted p-value < 0.05, Figure 6E).

## 4 Discussion

This study aimed to elucidate the mechanisms underlying the trade-off between grain weight and grain number in wheat, when grain weight is improved by genetic manipulations. To realize this objective, field evaluations of two genetically distinct wheat line groups, ExpA6 and GW2, were conducted under optimal conditions. The GW2 lines had 20 days longer crop cycle than the ExpA6 lines, but the climatic conditions during critical phenophases between both groups were similar across the two experimental years. As hypothesized, the transgenic and triple mutant lines displayed analogous phenology with their respective wild types, as well as plant height and architecture (data not shown).

Significant increments in grain weight were observed in the manipulated lines over the WTs, although differing in magnitude, i.e. by 8.6 and 23.7% in the *Ta*ExpA6 and *Ta*GW2 lines, respectively, without and with trade-off with grain number. These results agree with previous evaluations of the *Ta*ExpA6 and *Ta*GW2 lines (Wang et al., 2018; Zhai et al., 2018; Calderini et al., 2021), however, a direct comparison between both groups was feasible in our study as they were assessed in the same experiment sharing the same growing conditions and management. Notably, our study expands on previous work by demonstrating that both manipulations improved individual grain weight and grain dimensions across all grain positions of the spike, a feature not fully addressed in earlier research.

Comparative analyses between genetic resources and elite wheat varieties by Philipp et al. (2018) revealed that breeding process in wheat uniformly increased grain number and yield across the spike without altering individual spikelets relative contribution to overall yield. The observed increase in individual grain weight in our study aligns with these breeding trends, suggesting a similar pattern when grain weight is genetically improved. However, manipulation of *Ta*ExpA6 showed a higher impact on distal grains (G3 and G4) than on proximal ones (G1 and G2), while the TaGW2 triple mutant line showed similar increase across these grain positions.

Both genetic manipulations in our study successfully increased individual grain weight through an enhanced grain filling rate, maintaining the same grain filling duration as their respective WTs. Additionally, both modified lines reached higher maximum grain water content than their respective WTs, which has been ascribed as a driver of grain weight potential (Saini and Westgate, 1999; Pepler et al., 2006; Hasan et al., 2011; Alvarez Prado et al., 2013), suggesting increased sink strength which sequentially resulted in a higher grain filling rate and final grain weight. However, the impact of *Ta*ExpA6 and *Ta*GW2 manipulations on grain dimensions varied, since the transgenic approach primarily augmented grain length, while the triple mutation affected both length and width. This differential impact suggests distinct underlying mechanisms between manipulations. In the *Ta*ExpA6 line, the cloned expansin possibly facilitates cell wall loosening along the grains longitudinal axis, consistent with expansins mode of action (Cosgrove, 2023). Additionally, considering the cessation of cell proliferation in grains outer layers and starchy endosperm by 6 and 14 DAA, respectively (Olsen et al., 1992; Drea et al., 2005; Radchuk et al., 2011), the influence of the *Ta*ExpA6 gene seems more related to cell size than number.

Conversely, the *Ta*GW2 triple mutation appears to modulate both cell division and enlargement in maternal tissues around anthesis, resulting in longer and wider grains, in alignment with previous observations (Simmonds et al., 2016; Geng et al., 2017; Zhang et al., 2018). The significance of cell number and size in the outer tissues for potential grain weight previously exposed in bread wheat and across wheat ploidies (Lizana et al., 2010; Muñoz and Calderini, 2015; Brinton et al., 2017; Brinton and Uauy, 2019; Liu et al., 2020; Guo et al., 2022; Tillett et al., 2022), together with the findings from this study, support the hypothesis that potential grain weight is constrained by physical limitations imposed by maternal outer layers.

The effect of the manipulated lines on grain number per spike was contrasting as expected. The *Ta*ExpA6 gene did not affect this yield component, whereas the *Ta*GW2 triple mutation reduced it by 11.3% relative to the WT, across both seasons. This trade-off between grain weight and grain number when grain weight is increased has been widely reported (Wiersma et al., 2001; Brinton et al., 2017; Quintero et al., 2018; Wang et al., 2018; Golan et al., 2019; Milner et al., 2021). Notably, the magnitude of average grain weight increase did not directly correlate with grain number reduction, refuting the notion of a strict negative compensation between these spike yield components. These results align with the consensus that wheat grain filling is not source-limited (Slafer and Savin, 1994; Borrás et al., 2004; Reynolds et al., 2009; Quintero et al., 2018; Murchie et al., 2023; Slafer et al., 2023). Interestingly, the lack of effect on grain number showed by the ectopic expression of the *Ta*ExpA6 gene is consistent with two previous experiments at different plant rates (Calderini et al., 2021).

The *Ta*ExpA6 and *Ta*GW2 genes expression profile along with grain weight dynamics of the transgenic and mutant lines, allowed us to highlight a distinctive pattern in the effect of these manipulations on early grain development. The disruption of *Ta*GW2 gene in the triple mutant line improved grain weight above the WT from the onset of the grain filling period. In contrast, the higher grain weight reached by the *Ta*ExpA6 transgenic line became apparent at 21 DAA. Furthermore, the triple mutant exhibited heavier ovaries at pollination than the WT across different positions in the spikelet, indicating that the *Ta*GW2 gene functions in floret tissues before the gynoecium becomes a grain. This finding is in agreement with field experiments carried out by Simmonds et al. (2016), who identified increased carpel size and weight around anthesis as drivers of grain weight increase in *Ta*GW2 knockout mutants. In contrast, ovary weight at pollination did not differ significantly between *Ta*ExpA6 line and its WT in the present study, reinforcing the post-anthesis timing of the transgene effect as previously reported (Calderini et al., 2021).

It has been demonstrated that grain number and weight determinations overlap between booting and a week after anthesis (Calderini et al., 1999; Calderini et al., 2001; Ugarte et al., 2007; Hasan et al., 2011; Simmonds et al., 2014; Parent et al., 2017; Calderini et al., 2021). This overlap was proposed as the cause of the trade-off between this key yield components in wheat (Calderini et al., 2021). The effectiveness of the *Ta*ExpA6 gene in increasing grain weight without a trade-off with grain number supports this hypothesis, given that *Ta*ExpA6 expression begins at post-anthesis, particularly from 10 DAA onwards, and its effect on grain length and weight over the WT becomes detectable after 15 DAA. Conversely, when the enhancement commences at pre-anthesis as in *Ta*GW2 triple mutant, showed by both increased ovary size and grain weight, a trade-off occurs. Thus, our results suggest that the trade-off between grain weight and number is attributable to the temporal overlap of determinant periods for these grain components.

In addition to the effects of *Ta*ExpA6 and *Ta*GW2 gene manipulations on grain weight and number, recent elucidation of the role of the *GNI1* gene offers additional insights into the genetic regulation of these yield components. Research by Sakuma et al. (2019) highlighted that the *GNI1* gene, coding for a homeodomain leucine zipper class I transcription factor, plays a pivotal role in floret fertility in wheat. Their findings revealed an evolutionary adaptation in the expression of *GNI1*, leading to an increase in grain number per spikelet in domesticated wheat, suggesting a genetic basis for the observed trade-off between grain weight and grain number. Complementing this, Golan et al. (2019) proposed the *GNI-A1* gene, a variant of *GNI1*, as a mediator of this trade-off. Remarkably, *GNI1* is predominantly expressed in immature spikes before anthesis (Sakuma et al., 2019), further highlighting the relevance of the overlap between grain number and weight determinations as key to unravel the underlying causes for the trade-off. Additionally, Xie and Sparkes (2021) found an overlapping between major regions associated with grain number and grain weight in a mapping population of 226 RILs .

The integration of these insights with the results of our study suggests a multifaceted genetic network influencing wheat yield. While our study focuses on the phenotypic outcomes of *Ta*ExpA6 and *Ta*GW2 gene manipulations, the role of the *GNI1* gene underscores the importance of genetic control in floret fertility and assimilate allocation. This understanding complements our observations of the trade-off between grain weight and grain number, indicating that both targeted genetic modifications and natural genetic variations, such as those in *GNI1*, are crucial in determining these key agronomic traits. Therefore, a comprehensive strategy that considers both natural gene variants like *GNI1* and targeted modifications such as *Ta*ExpA6 and *Ta*GW2 may be essential for the development of wheat varieties that optimally balance grain weight and number.

## 5 Concluding remarks

In conclusion, the results of this study emphasize the complexity of the genetic control of grain weight and number in wheat. The findings reveal that modification of the *Ta*ExpA6 gene enhances grain weight without reducing grain number, indicating a deviation from the traditionally assumed trade-off between these yield components. In contrast, alterations in the *Ta*GW2 gene, while increasing grain weight, also result in a reduction in grain number, aligning with the conventional understanding of this trade-off. These outcomes highlight distinct genetic pathways influencing wheat yield traits. The differential impacts of *Ta*ExpA6 and *Ta*GW2 on wheat grain development, and the overlapping of both yield components determination between booting and a week after anthesis, offer valuable insights for future wheat breeding programs aimed at improving yield through targeted genetic modifications.

## 6 Data availability statement

Original contributions and data presented in this study is available within the article or its supplementary materials. Further inquiries can be directed to the corresponding author.

## 7 Author contributions

DFC conceived the project and coordinated the experiments and data analysis; JC proposed the molecular analysis and evaluated molecular data; LV conducted field experiments and data analyses. LV and DFC wrote the manuscript with contributions from JC.

## Supporting information

Trade-off between grain weight and grain number in wheat

## 9 Acknowledgements

The authors wish to warmly thank Dr. Emma Wallington and the National Institute of Agricultural Botany (NIAB, UK) for kindly facilitating the ExpA6 lines, and Prof. Cristobal Uauy and John Innes Center (JIC, UK) for their generous provision of the GW2 lines. We also thank the experimental field staff of the Austral Farming Experimental Station of Universidad Austral de Chile for the technical assistance provided. Technical contributions and management of field trials by Marcelo Castro Moraga and Cristóbal Castro Villalón is truly appreciated. We are also very grateful to María Beatriz Ugalde Jaramillo and Dr. Anita Arenas Miranda (Plant Nutrition and Genomics Lab – UACh) for performing the molecular analysis included in this study. The samples processing by Beatriz Shibar (UACh) and Magda Lobnik is recognized. J.C. was supported by ANID— Millennium Science Initiative Program—ICN17-022 and FONDECYT Regular 1230833. L.V. was supported by PhD scholarship 21220957-2022 from ANID, Chile.

