## Supplementary material for "The trade-off between grain weight and grain number in wheat is explained by the overlapping of the key phases determining these major yield components": Trade-off between grain weight and grain number in wheat

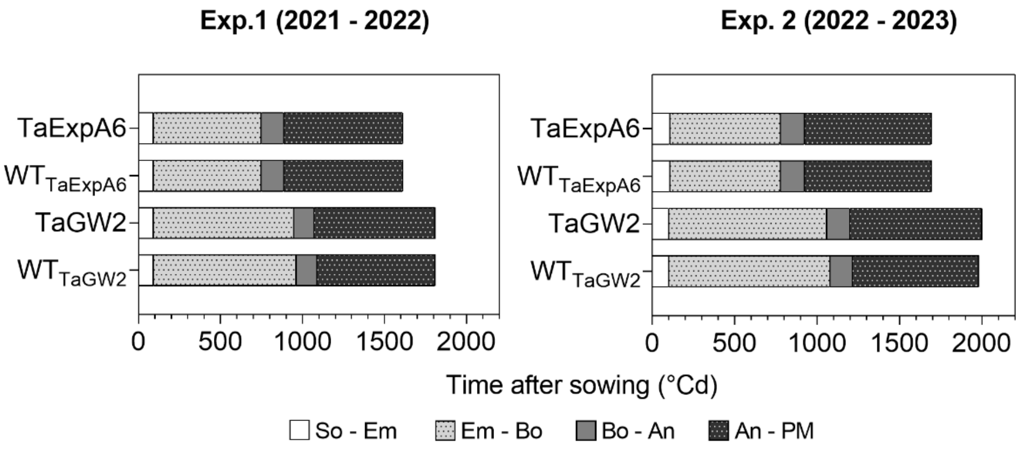


**Figure S1.** Phenological phases of *Ta*ExpA6 and *Ta*GW2 lines, and their respective wild types (WT), from sowing to physiological maturity in Experiments 1 and 2. Bars show the duration of each phase in thermal time: from sowing to seedling emergence (So-Em), from seedling emergence to booting (Em-Bo), from booting to anthesis (Bo-An) and anthesis to physiological maturity (An-PM).

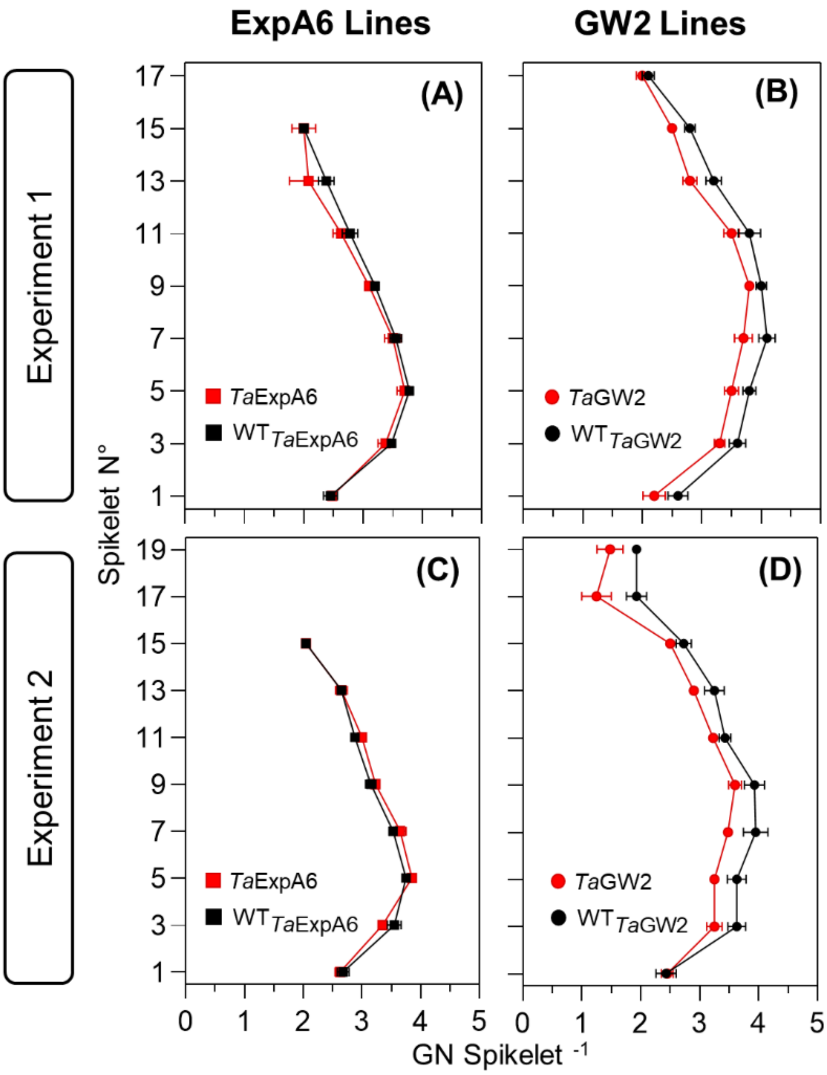


**Figure S2.** Grain number per spikelet along the spike of ExpA6 lines (A, C) and GW2 lines (B, D) in experiments 1 (upper panel) and 2 (lower panel). *Ta*ExpA6 line and the triple mutant of *Ta*GW2 gene, and their WTs are denoted by red and black symbols, respectively.

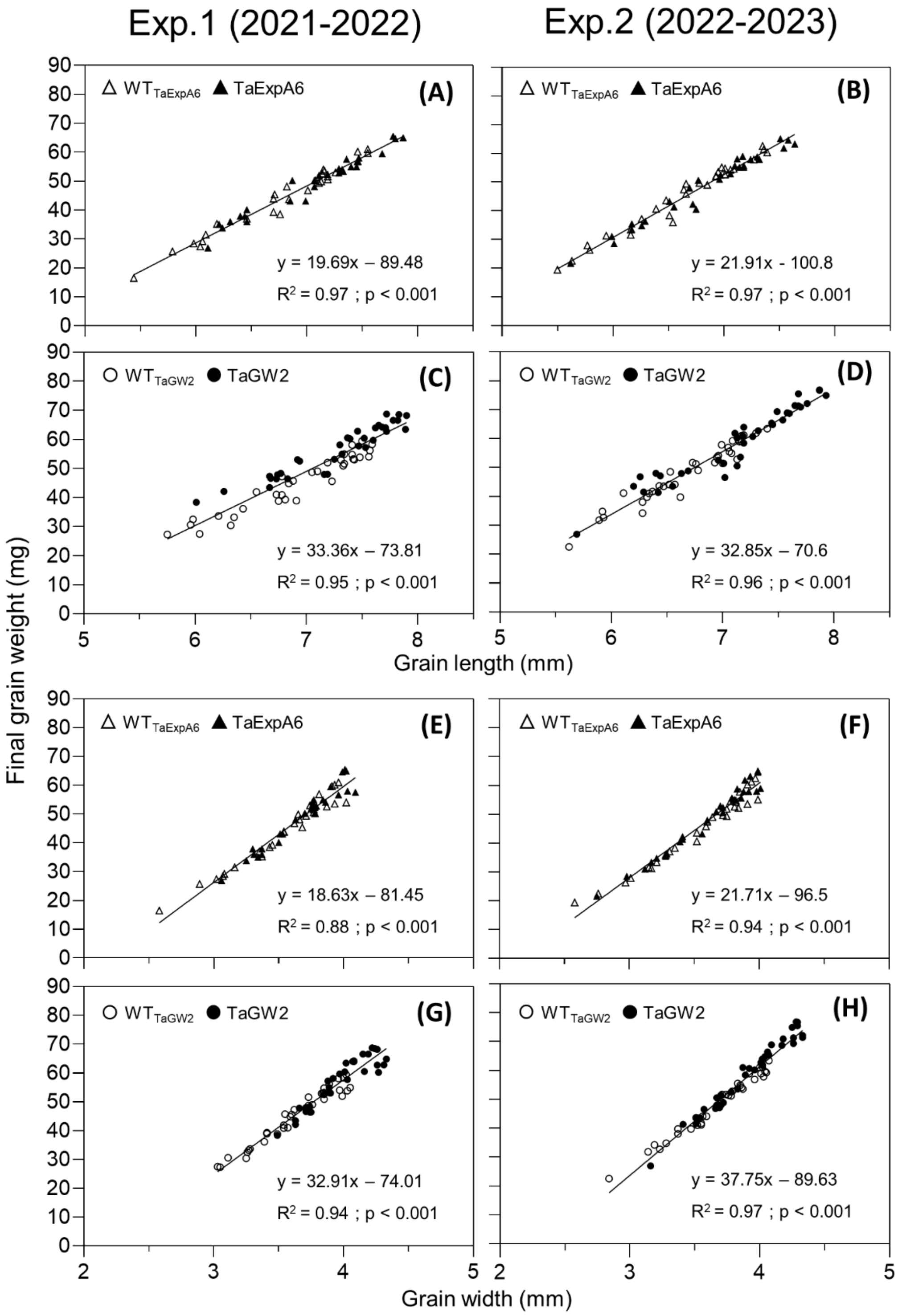


**Figure S3.** Relationship between grain weight and grain dimensions of grain positions G1, G2, G3 and G4 from each spikelet along the spike in Experiments 1 (left panel) and 2 (right panel). **(A, B)** Relationship between grain weight and grain length in ExpA6 lines and **(C, D)** GW2 lines. **(E, F)** Relationship between grain weight and grain width in ExpA6 lines and **(G, H)** GW2 lines.

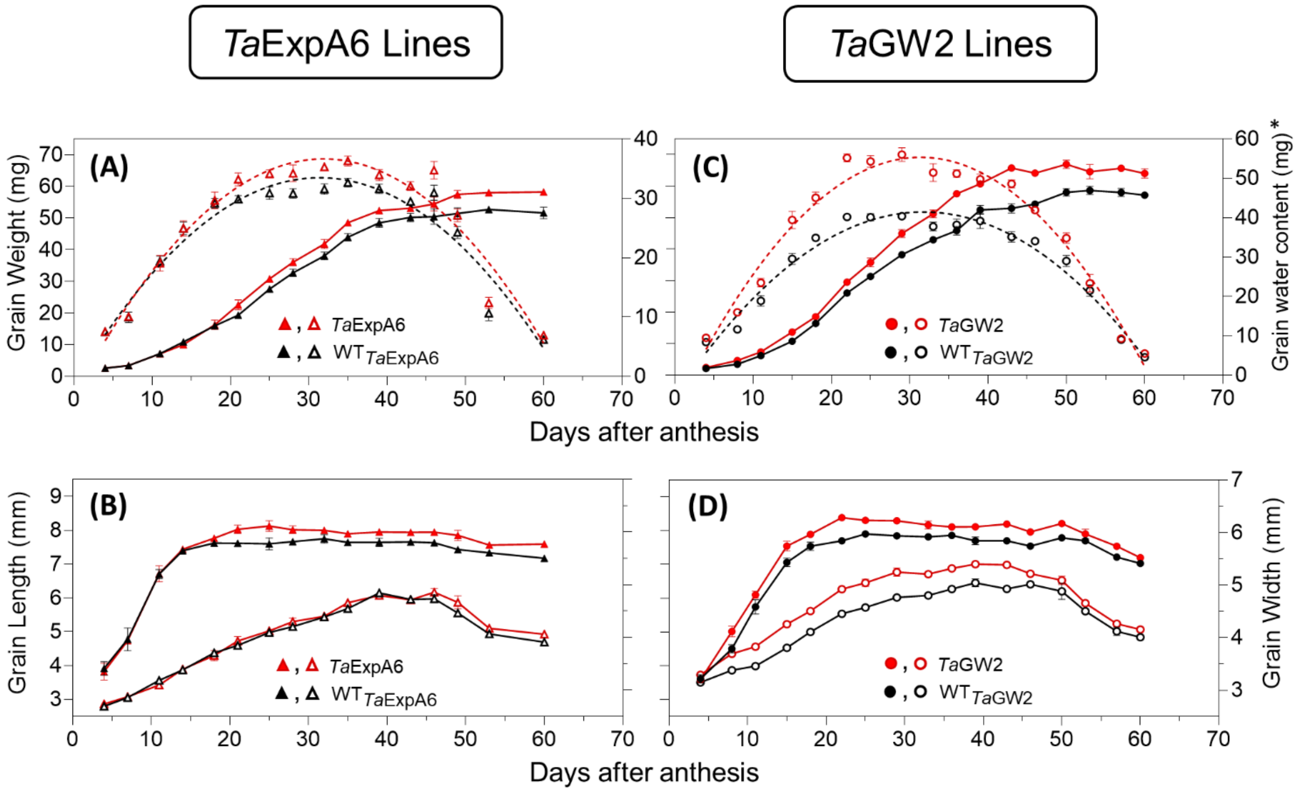


**Figure S4.** Grain weight dynamics and gene expression. **(A)** Grain weight and water content dynamics at grain position G3 from the central spikelets of the spike corresponding to ExpA6 lines and **(C)** GW2 lines. **(B)** Grain length and grain width dynamics at the same grain position corresponding to ExpA6 lines and **(D)** GW2 lines. **Note that different scales were used to plot grain water content of ExpA6 and GW2 lines.*

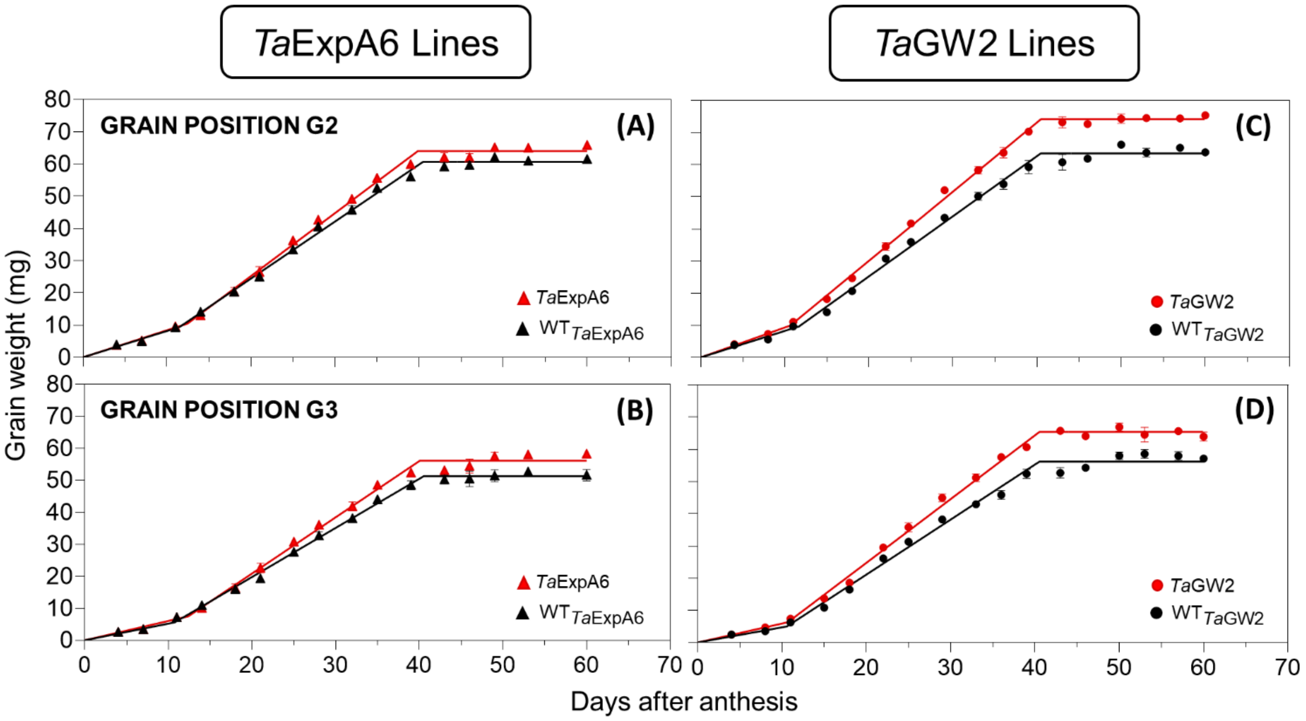


**Figure S5.** Tri-linear model fitted to grain weight dynamics at grain positions G2 (upper panel) and G3 (lower panel) in ExpA6 **(A, B)** and GW2 **(C, D)** lines.

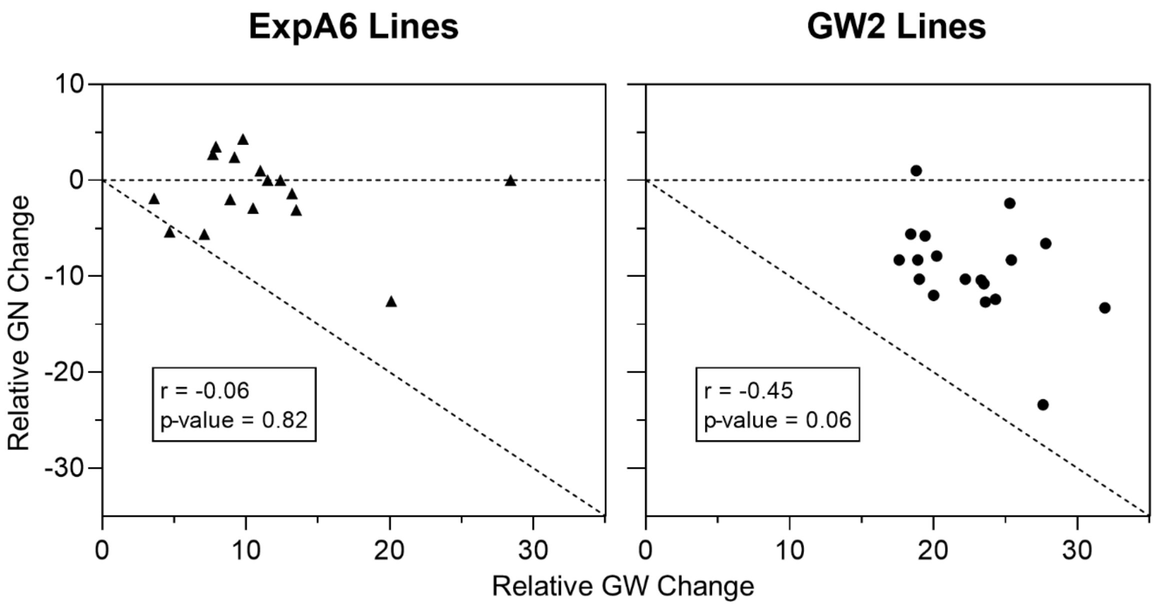


**Figure S6.** Relationship between relative average increase in grain weight in each spikelet along the spike and the relative change in grain number per spikelet. Horizontal dashed line represents no change in grain number, while diagonal dashed line represents a proportional inverse relationship between grain weight and grain number.

**Table S1.** Equation parameters of the tri-linear model fitted in the individual seed weight dynamics of grain positions G2 and G3

| Grain position | Group | Line | $R^2$ | y1 (Filling rate; lag phase) | y2 (Filling rate; linear phase) | X1 (duration of lag phase; days) | X2 (duration of grain filling; days) | Plateau (Stabilised grain dry weight; mg) |
|---|---|---|---|---|---|---|---|---|
| G2 | *Ta*ExpA6 | $WT_{TaExpA6}$ | 0.99 | 0.82 | 1.77 | 11.5 | 40.5 | 60.7 |
| | | TaExpA6 | 0.99 | 0.84 | 1.95 | 12.4 | 39.9 | 64.0 |
| | | p-value | - | ns | * | ns | ns | - |
| | *Ta*GW2 | $WT_{TaGW2}$ | 0.99 | 0.83 | 1.86 | 11.6 | 40.6 | 63.6 |
| | | TaGW2 | 0.99 | 0.93 | 2.16 | 10.7 | 40.3 | 74.1 |
| | | p-value | - | ns | ** | ns | ns | - |
| G3 | *Ta*ExpA6 | $WT_{TaExpA6}$ | 0.98 | 0.52 | 1.53 | 10.7 | 40.5 | 51.3 |
| | | TaExpA6 | 0.99 | 0.60 | 1.76 | 12.4 | 40.1 | 56.1 |
| | | p-value | - | ns | ** | ns | ns | - |
| | *Ta*GW2 | $WT_{TaGW2}$ | 0.99 | 0.47 | 1.68 | 10.3 | 41.1 | 56.4 |
| | | TaGW2 | 0.99 | 0.59 | 2.01 | 10.9 | 40.1 | 65.2 |
| | | p-value | - | ns | ** | ns | ns | - |

Extra sum of squares F test was used for the comparison of the equation parameters between each modified line and its respective WT. Different letters indicate significant effects: *, $P < 0.05$;**, $P < 0.01$; ns, not significant.
